# Identification of enhancer chromatin signatures involved in dopaminergic induction through multi-omics analysis

**DOI:** 10.1101/2025.08.14.670306

**Authors:** Mayela Giacoman-Lozano, César Meléndez-Ramírez, Adolfo López-Ornelas, Aylin Del Moral-Morales, Rosa Gloria Rebollar-Vega, Ernesto Soto-Reyes, Iván Velasco, Raquel Cuevas-Diaz Duran

## Abstract

The establishment of precise transcriptional programs during neuronal development depends on a complex and dynamic landscape of regulatory elements. Here, we combined histone modification ChIP-seq, chromatin accessibility profiling, and transcriptomics to map active enhancer repertoires during the differentiation of human embryonic stem cells into midbrain dopaminergic neurons. Our integrative analysis revealed thousands of stage-specific enhancers, over half of which were previously unannotated, and uncovered coordinated chromatin and transcriptional transitions linking pluripotency exit to neuronal lineage commitment. Functional enrichment of enhancer-linked target genes delineated distinct regulatory programs, pluripotency maintenance in undifferentiated cells and midbrain specification in differentiated neuron, while transcription factor motif analysis identified regulatory modules, including a dopaminergic-specific RFX4 network. We further characterized enhancer-gene relationships showing concordant changes in chromatin state and expression for neuronal genes, suggesting enhancer-driven control of dopaminergic identity. These findings provide a genome-scale framework for interpreting how dynamic enhancer landscapes encode developmental fate decisions in the human nervous system.

## Background

Stem cell differentiation is orchestrated by epigenetic mechanisms that regulate dynamic gene expression in each stage of development. Genomic enhancers are regulatory regions that play a crucial role during the differentiation process by interacting with promoter regions to increase transcription likelihood. In contrast to promoters, enhancers do not interact solely with genes in close proximity (1). Additionally, their activity is highly cell-type and developmental stage specific (2), making them more elusive and their identification more challenging. Putative enhancers have been identified using a wide variety of sequencing methods. Genome-wide sequencing technologies have allowed for genome-wide maps of DNA-interacting proteins associated with enhancer activity. For example, chromatin immunoprecipitation followed by high throughput sequencing (ChIP-Seq) is used to identify the post-translational histone modifications H3K27ac and H3K4me1 as well as transcription factor EP300 enrichment (3). Open-chromatin profiles as well as H3K4me1 and H3K27ac histone modification enrichment are highly correlated with enhancer activity. Since H3K4me1 alone can mark both poised and active enhancers, while H3K27ac is associated with active enhancers and promoters, their combined analysis allows for a more refined identification of putative regulatory elements (4).

Given the highly dynamic nature of enhancer activity across different cell types and developmental stages, identifying key regulatory elements involved in neuronal differentiation requires a comprehensive, integrative approach. Neurons are among the cell types with the largest number of specific enhancers, whose activity is variable between different developmental stages (5) and different neuronal types (6). Generation of human dopaminergic neurons (mDA) from human embryonic stem cells has been achieved (7). These differentiation models have emerged as useful tools for the study and annotation of key genome regulatory regions during development (8). This differentiation model results in in vitro differentiated neurons that express dopaminergic markers, such as TH, GIRK2, LMX1A, EN1 and FOXA2. Also, these neurons have electrophysiological activity that resembles the response of mDA. These cells are of clinical interest due to their role in the pathophysiology of Parkinson’s Disease. Using a variety of sequencing techniques simultaneously provides information about the association between chromatin structure, biochemical annotations, and gene expression. Integration of these assays during dopaminergic differentiation provides useful information about regulatory regions involved in neural development and dysfunction (9).

Although the significance of enhancers in the development of the nervous system is well-recognized, the specific regulatory elements that guide dopaminergic differentiation are still not fully understood. A more thorough identification of the active enhancers involved in this process could enhance our understanding of the transcriptional mechanisms that dictate neural fate choices and regulate genes linked to diseases. This study aims to identify and characterize active enhancers during dopaminergic differentiation by integrating multiple sequencing datasets. ChIP-Seq for H3K4me1 and H3K27ac histone modifications was performed during embryonic stem cell differentiation to dopaminergic neurons. For an integrated approach, data obtained from these experiments as well as ATAC-Seq and RNA-Seq datasets previously performed on the same differentiation model were analyzed to select enhancer candidates for further validation.

## Materials and Methods

### hESCs culture and dopaminergic differentiation

H9 human embryonic stem cells expressing GFP (H9GFP) (Domingo-Reines et al., 2017), were defined as day zero (D0) for this analysis. We then differentiated this cell line into dopaminergic neurons following the dual-SMAD inhibition and floor-plate induction protocol described by Kriks et al. (2011), with minor modifications. Differentiation continued until day 28, when chromatin was extracted for analysis.

### Chromatin extraction and ChIP-Seq

Chromatin was extracted from cells at the pluripotent stage (D0) and after 28 days of dopaminergic differentiation. Cells were cross-linked with 1% formaldehyde for 10 min, quenched with 1.38 M glycine, and washed with cold PBS. After removing PBS, cells were lysed in SDS-based buffer with protease inhibitors, incubated at 4°C, and collected using a cell scraper. Chromatin was sheared using a Covaris S200 sonicator for 80 seconds. DNA fragmentation (200–500 bp) was assessed by de-crosslinking, RNA digestion, phenol-chloroform extraction, and agarose gel electrophoresis. ChIP was performed with the One-day ChIP kit (Diagenode) using 1–2 million cells per experiment and 200,000 cells as input. The number of cells was estimated through histone quantification using Bradford protein assay to avoid lifting cells for counting before ChIP. H9 ESCs were used as reference for protein quantification. Antibodies against H3K27ac (Active Motif) and H3K4me1 (Diagenode) were used, and ChIP efficiency was validated by ChIP-PCR before sequencing. Immunoprecipitated DNA was purified using the ChIP Elute Kit and quantified with a Qubit Fluorometer to determine library preparation cycles. Samples were either stored at −20°C or processed immediately.

### ChIP-Seq data analysis

ChIP-Seq libraries were sequenced on an Illumina NovaSeq 6000 sequencer to obtain 100-bp single-end reads. Quality control of sequenced reads was performed using FastQC [FastQC: A quality control tool for high throughput sequence data. Available at: http://www.bioinformatics.babraham.ac.uk/projects/fastqc/ (Accessed: June 2024).]. Reads were aligned to the GRCh38/hg38 human reference genome using Bowtie2 v2.4.4 [https://doi.org/10.1038/nmeth.1923] with default parameters. Duplicated reads were located and tagged using Picard’s MarkDuplicates tool v2.27.5 [http://broadinstitute.github.io/picard]. Subsequently, reads were filtered using Samtools v1.13 [https://doi.org/10.1093/gigascience/giab008] to exclude those flagged as PCR duplicates, unmapped, non-primary alignments, multi-mapping, or reads failing platform quality checks. Furthermore, reads with mapping quality MAPQ < 30 and those mapped to mitochondrial DNA were also removed from downstream analysis. After post-alignment filtering, peaks were called with MACS2 v2.2.7.1 (10) callpeak function with modified parameters (--broad --nomodel --extsize) and a q-value cutoff of 0.1. Extension sizes 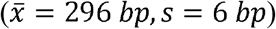 were obtained by previously running macs2 callpeak function and allowing it to build the shifting model. For signal track visualization, normalized fold enrichment tracks were generated using macs2 callpeak function with --SPMR parameter and then passing the bedgraph outputs to the macs2 bdgcmp function with -m FE setting. Resulting peaks overlapping ENCODE blacklisted regions (11) were filtered out and the final list of peaks representing each histone modification and time point is included in Additional File 1.

### Peak annotation and correlation to gene expression

For each histone modification, resulting peaks were annotated using the annotatePeaks function included in Homer (12). We used the distance to the nearest TSS, the nearest neighbor and its gene type to evaluate correlation between histone enrichment and gene expression. We used RNA-seq datasets previously published by our lab (13) to obtain the mean log2-transformed FPKM values of protein-coding genes and we defined three gene expression levels: low (log2FPKM < 4), medium (4 <= log2FPKM < 8), and high (log2FPKM >=8). Histone modification enrichment profiles around the TSS (±5*kb*) were generated for each level of gene expression (Figure 1E). Annotated peaks are included in Additional File 2.

**Figure 1.**
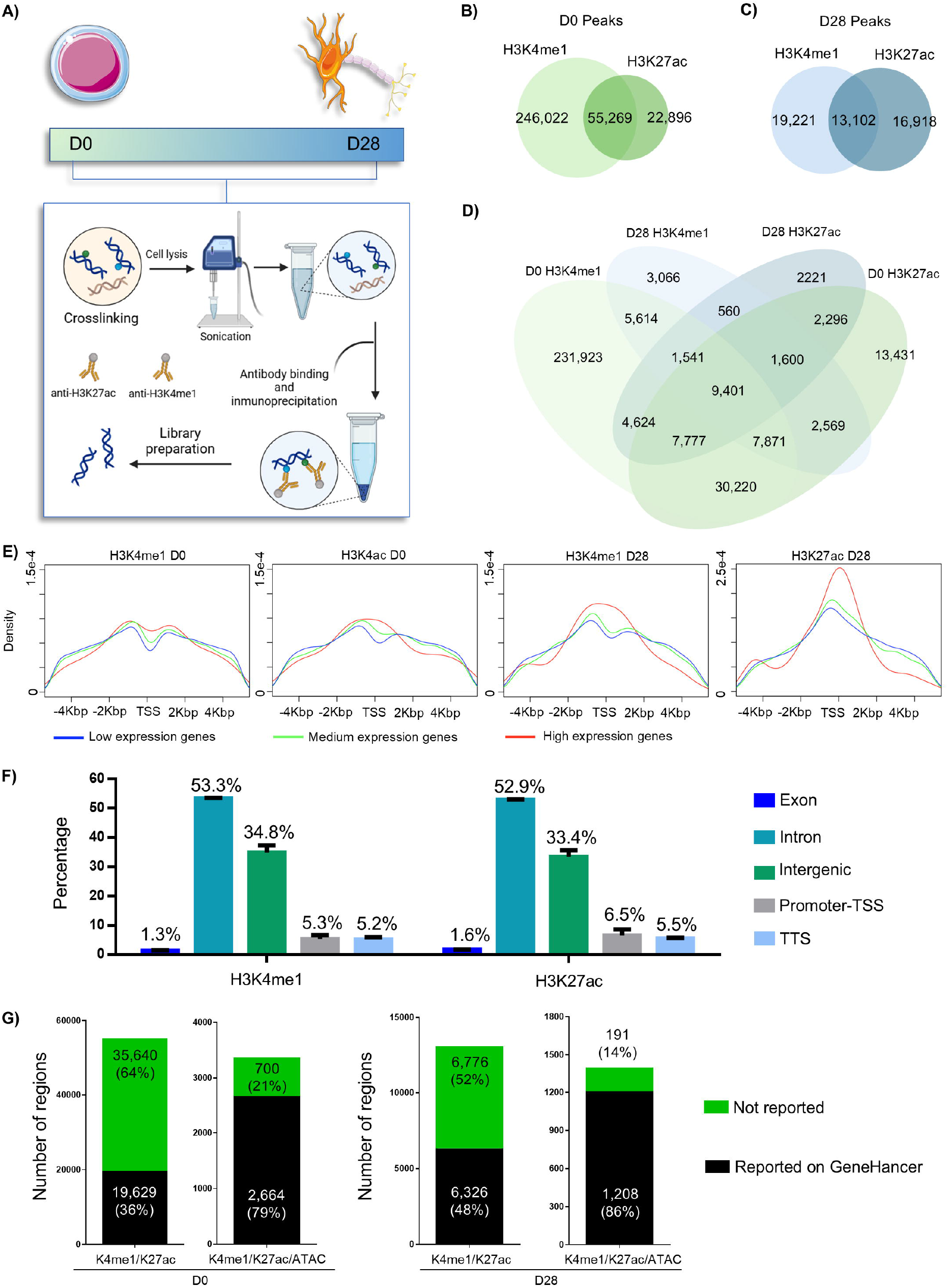
Genome-wide identification of putative enhancer regions in hESC and mDA cells using histone modification ChIP-seq. **(A)** Schematic representation of the ChIP-seq workflow: hESC (D0) and midbrain mDA differentiated from hESC (D28) were crosslinked, lysed, and subjected to chromatin fragmentation by sonication. ChIP was performed using antibodies against H3K4me1 and H3K27ac. Purified DNA was used for library preparation and sequencing; two biological replicates were pooled for each condition. **(B)** Overlap of significant peaks detected for each histone modification at D0. **(C)** Overlap of significant peaks detected for each histone modification at D28. **(D)** Venn diagram showing the global overlap of significant peaks of both histone modifications between D0 and D28. **(E)** Meta-analysis of histone modifications: H3K4me1, H3K4ac (D0), and H3K4me1, H3K27ac (D28) enrichment at ±4 kb around transcription start sites (TSS) of genes with low (blue), medium (green), and high (red) expression levels. In both time points, higher enrichment of active histone modifications is observed at promoters of highly expressed genes. **(F)** Average genomic distribution of H3K4me1- and H3K27ac-enriched regions in both evaluated days. Percentages indicate the proportion of peaks in each genomic category. **(G)** Bar plots showing the number of genomic regions marked by H3K4me1/K27ac that are already reported in the GeneHancer database; from left to right: D0 without chromatin accessibility (ATAC-seq overlap), D0 with ATAC-Seq overlap, D28 without ATAC-Seq overlap, and D28 with ATAC-Seq overlap. Black bars indicate reported regions, while green bars represent regions not previously reported. Percentages correspond to the proportion of regions within each category.

### Chromatin state comparisons with H3K27ac epigenomic profiles of related cell-types

To compare the chromatin states between other cell-types, we downloaded H3K27ac datasets derived from neural precursor cells (NPC), and specialized neuronal subtypes including motor, glutamatergic, and gabaergic neurons. NPC and motor neuron H3K27ac profiles were obtained from the ENCODE portal (14) (http://www.encodeproject.org) with the following identifiers: ENCSR449AXO and ENCSR489LNU. Glutamatergic and gabaergic neuron H3K27ac profiles were retrieved from Gene Expression Omnibus (https://www.ncbi.nlm.nih.gov/geo/) with GSM5862749 and GSM5862738 accession numbers. We performed H3K27ac profile comparison using Jaccard index and hierarchical clustering.

### Identification and clustering of active enhancers

For each time point, we defined active enhancers as those regions enriched in both histone modifications (H3K4me1 and H3K27ac) and chromatin accessibility (ATAC-seq). We used previously published chromatin accessibility profiles found in dopaminergic induction (13). Overlap of marks in each time point were obtained using bedtools v2.31.0 intersect function (15) (Additional File 3). Then, a list of consensus putative enhancers were derived by merging the regions belonging to each time point using bedtools v2.31.0 merge function (15). These consensus regions were considered as pseudo-genes and the number of read counts mapped to them was obtained by implementing bedtools v2.31.0 multicov function (15) with alignment files.

To compare the enrichment of histone modifications and chromatin accessibility profiles between the two time points (D0 vs D28), we performed a differential enrichment analysis using consensus regions’ counts with DESeq2 v1.38.3 (16). Regions with log2-transformed fold-enrichment greater than 1.5 and *p-value* less than 0.05 were considered differentially enriched. Hierarchical clustering of these regions was performed using Ward’s method based on the Euclidean distance matrix of z-score scaled normalized counts and depicted in a heatmap (Figure 3A). To increase the reliability of the active enhancer regions inferred, we performed an overlap between differentially enriched regions and enhancer regions annotated in the GeneHancer database included in GeneCards Suite v4.14 (17).

### Identification of TF binding in active enhancer regions

To identify TFs potentially binding active enhancers overlapping GeneHancer regulatory regions, we performed a motif analysis using ENCODE-motifs (18) downloaded from https://compbio.mit.edu/encode-motifs/. Active enhancer region sequences were downloaded from the hg38 reference genome using the BSgenome R library [doi:10.18129/B9.bioc.BSgenome]. For each cluster identified, motifs were used to scan active enhancer regions using FIMO (19) with a False Discovery Rate (FDR) < 0.05.

## Results

### ChIP-seq histone marks delimit common and specific putative enhancers

To identify genome-wide putative enhancer regions, we performed a chromatin immunoprecipitation followed by sequencing (ChIP-seq) assay. We used antibodies against enhancer-associated histone H3 modifications, H3K4me1 and H3K27ac in a human embryonic stem cell line (hESC) and in early mDA differentiated from hESC using a 28-day previously reported floor-plate protocol (7)(13). hESC and mDA samples were identified as D0 and D28 according to induction day. By pooling two biological replicates, we obtained ChIP-seq samples for two time points and two H3 histone modifications (D0H3K4me1, D0H3K27ac, D28H3K4me1, and D28H4K27ac) following the methodology depicted in Figure 1A. Genome-wide, we identified 298,971 and 75,165 significant peaks in D0 samples corresponding to H3K4me1 and H3K27ac modifications respectively. As observed in Figure 1B, a total of 55,269 significant peaks found in D0 depicted both histone modifications, representing putative enhancers. Interestingly, D28 samples displayed a smaller number of significant peaks: 32,222 and 30,020 derived from H3K4me1 and H4K27ac modifications. Only 13,102 regions were characterized with both histone modifications (Figure 1C). In both time points, regions enriched in H3K4me1 but not in H3K27ac are considered inactive enhancers and those enriched in H3K27ac but not in H3Kme1 represent other active regions, for example, promoters. A list of resulting significant peaks for each histone modification and time point is included in Additional File 1.

We compared specific and overlapping histone modifications between time points through a Venn diagram (Figure 1D). A total of 9,401 regions were enriched in both histone modifications and in both D0 and D28 samples. These common enhancer regions potentially regulate housekeeping genes. Moreover, we found 30,220 and 560 regions with overlapping histone modifications exclusively in D0 and D28 samples respectively. These regions represent putative enhancer regions potentially regulate cellular functions specific to each time point. We also observed regions enriched only in H3K4me1 modification in both D0 and D28 time points, suggesting inactive enhancers. Similarly, 2,296 regions were enriched only in H3K27ac mark in both time points, suggesting promoter regions. A total of 231,923 and 3,066 regions depicted H3K4me1 enrichment without H3K27ac modification only in D0 or D28 correspondingly. Furthermore, 13,431 and 2,221 regions in D0 and D28 were enriched only in H3K27ac.

### Characterization of histone modification sites and correlation with gene expression

To study the distribution of histone modifications near transcription start sites (TSS), we generated composite genome-wide enrichment profiles of each mark in a range of ±5 *kb* from protein-coding genes’ TSS. Different profiles were generated according to genes’ expression levels obtained from previously published RNA-seq data sets (13) (Figure 1E). Profiles show that H3K27ac modifications in both D0 and D28 are enriched around the TSS, however this trend is more robustly depicted in highly expressed genes. In contrast, H3K4me1 enrichment is lower in the TSS and increases reaching a maximum approximately at ±1 *kb* depicting a bimodal distribution, however this trend is not observed in the profile of highly expressed genes of D28. These results suggest that during the differentiation process, the H3K4me1 signal is transitioning from a bimodal to a unimodal distribution allowing promoters to become poised, a fundamental lineage specification mechanism (20)(21). Moreover, in D28 H3K27ac histone modification enrichment profiles appear positively correlated with gene expression levels, and they are consistent with previously reported distribution patterns.

To analyze the genomic location of histone modification enrichment, we intersected peaks with different genomic region classes: exons of annotated genes, proximal promoter regions and TSS (1kb upstream and 100 bps downstream), transcription termination sites (TTS), intronic, and intergenic regions (Additional File 2). For both modifications, enrichment regions in D0 and D28 time points were mostly located in non-coding DNA (86-88%), namely introns and intergenic regions, suggesting putative enhancers. These results are consistent with previous reports where H3K4me1 is generally associated with distal cis-regulatory elements (22) and where H3K27ac distinguishes active enhancers from poised ones (4). The genomic distribution of both histone modifications did not show significant differences between time points, suggesting a stable distribution. Moreover, the distribution of both modifications is also very similar (Figure 1F), pinpointing at overlapping marks in non-coding regions, a well-known characteristic of active enhancers (23).

### Putative enhancers overlap with chromatin accessibility and GeneHancer annotated regions

Active enhancers are DNA regulatory elements that increase the transcription of target genes. Besides being enriched in H3K4me1 and H3K27ac modifications, active enhancers are also located within open chromatin regions (24)(25). To infer the proportion of putative enhancers (overlapping H3K4me1 and H3K27ac) that are active at each time point according to their chromatin accessibility, we used a previously published Assay for Transposase-Accessible Chromatin using sequencing (ATAC-seq) dataset (13). Furthermore, we also compared putative enhancer regions to the GeneHancer regulatory elements database (17), a comprehensive source of human enhancer and promoter regions derived from the integration of 9 databases. We found that 36% (19,629 out of 55,269) and 48% (6,326 out of 13,102) of the regions enriched in both H3K4me1 and H3K27ac marks in D0 and D28 respectively have evidence in GeneHancer database. However, only 6.1% (3,364 out of 55,269) and 10.7% (1,399 out of 13,102) of the regions with overlapping marks in D0 and D28 respectively, reside in open chromatin. Moreover, 79% (2,664 out of 3,364) and 86% (1,208 out of 1,399) of the regions depicting the three attributes (histone modifications and open chromatin) obtained from D0 and D28 samples are annotated in GeneHancer database, increasing the reliability of predicted active enhancers. Therefore, we defined active enhancers as those open chromatin regions enriched in both histone modifications. Bar plots depicting the frequency of histone modifications and chromatin accessibility regions overlapping GeneHancer elements for each time point are found in Figure 1G. A list of regions enriched in both histone modifications classified by chromatin accessibility and overlap with GeneHancer elements is included in Additional File 3.

### Time point-specific active enhancer regions are associated with pluripotency or neuronal genes

To evaluate the overlap of histone modifications with chromatin accessibility, we analyzed regions located within a vecinity of 20 Kb of NANOG and FOXA1, transcription factors known to be involved in pluripotency maintenance and dopaminergic neuron specification respectively. As observed in Figure 2A, two putative enhancers overlapping GeneHancer regions (GH12J007786, GH12J007796) surround NANOG gene, and they are both characterized by decreased chromatin accessibility in D28. Moreover, a decrease in H3K27ac enrichment from D0 to D28 is also observed in the enhancer spanning NANOG’s TTS (GH12J007796). These results suggest that NANOG’s transcriptional activity is potentially regulated by these active enhancers. Notably, enrichment of H3K27ac modification is only observed in neural precursor cells, whereas it is low or absent in glutamatergic neurons and motor neurons.

**Figure 2.**
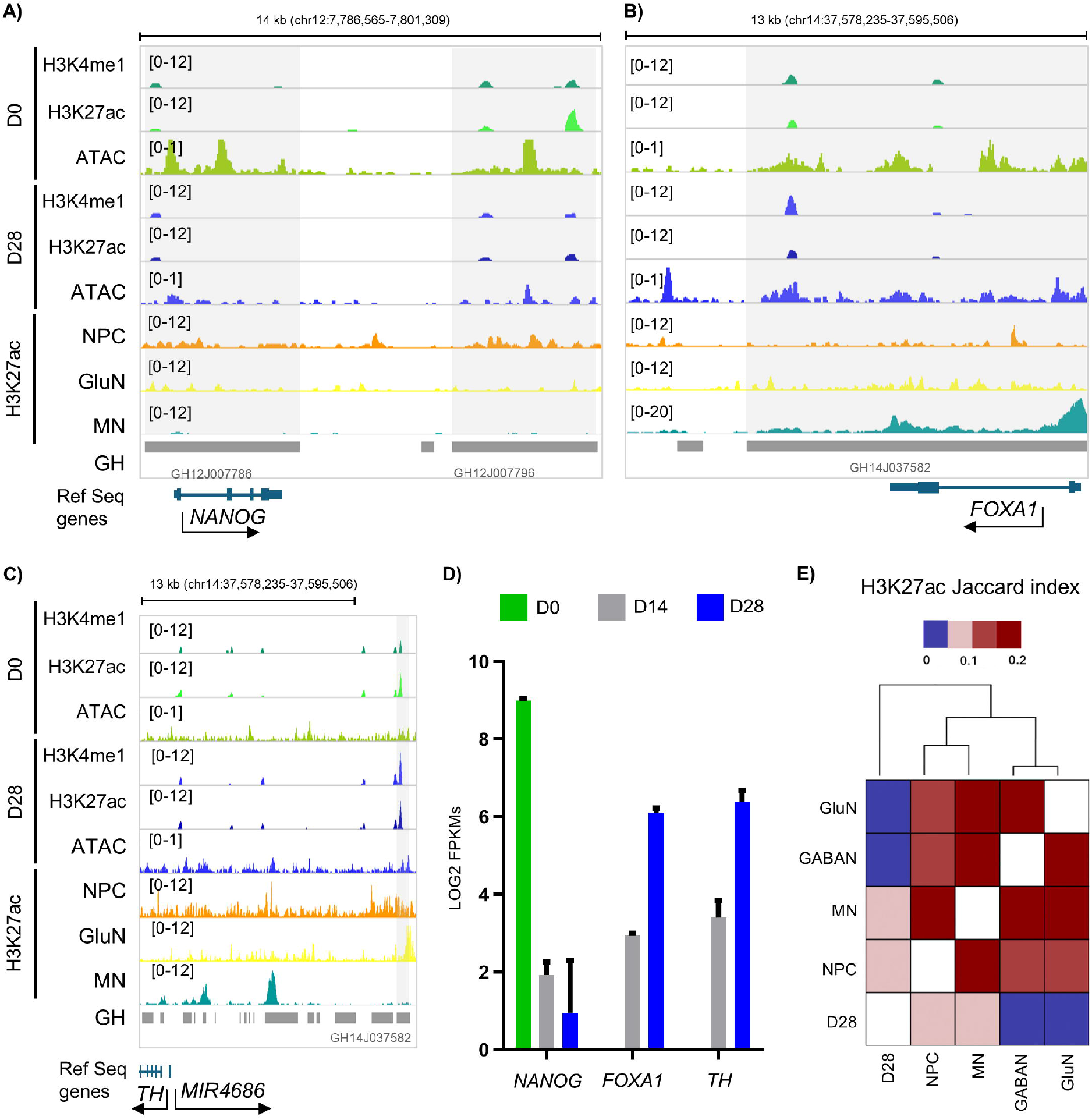
Examples of time point–specific active enhancers associated with pluripotency- and neuron-related genes. **(A)** Genome browser tracks showing ChIP-seq enrichment for H3K4me1 and H3K27ac at two putative enhancer regions flanking the pluripotency gene NANOG. **(B)** Putative enhancer that is located downstream of the FOXA1 transcription termination site (TTS) that overlaps a GeneHancer locus. **(C)** Putative enhancer located ~40.7 kb upstream of the TH transcription start site (TSS), identified in the GeneHancer database (GH11J002212). This region maintains H3K27ac enrichment across time points but gains additional H3K4me1 in D28 Tracks display H3K27ac, H3K4me1, and ATAC-seq profiles in D0 and D28 samples, along with H3K27ac profiles from neural progenitor cells (NPC), glutamatergic neurons (GluN), motor neurons (MN), and GeneHancer annotations (GH). Signal intensity ranges are indicated in brackets to the right of each track. **(D)** Bar plots showing LOG2 FPKMs for NANOG, FOXA1, and TH transcripts across the dopaminergic differentiation. Temporal patterns of chromatin accessibility are correlated with transcriptomic profiles. **(E)** Heatmap showing pairwise Jaccard similarity indices for H3K27ac-enriched regions between D28, neural progenitor cells (NPC), motor neurons (MN), GABAergic neurons (GABAN), and glutamatergic neurons (GluN). Higher similarity values (red) indicate greater overlap in active enhancer profiles.

**Figure 3.**
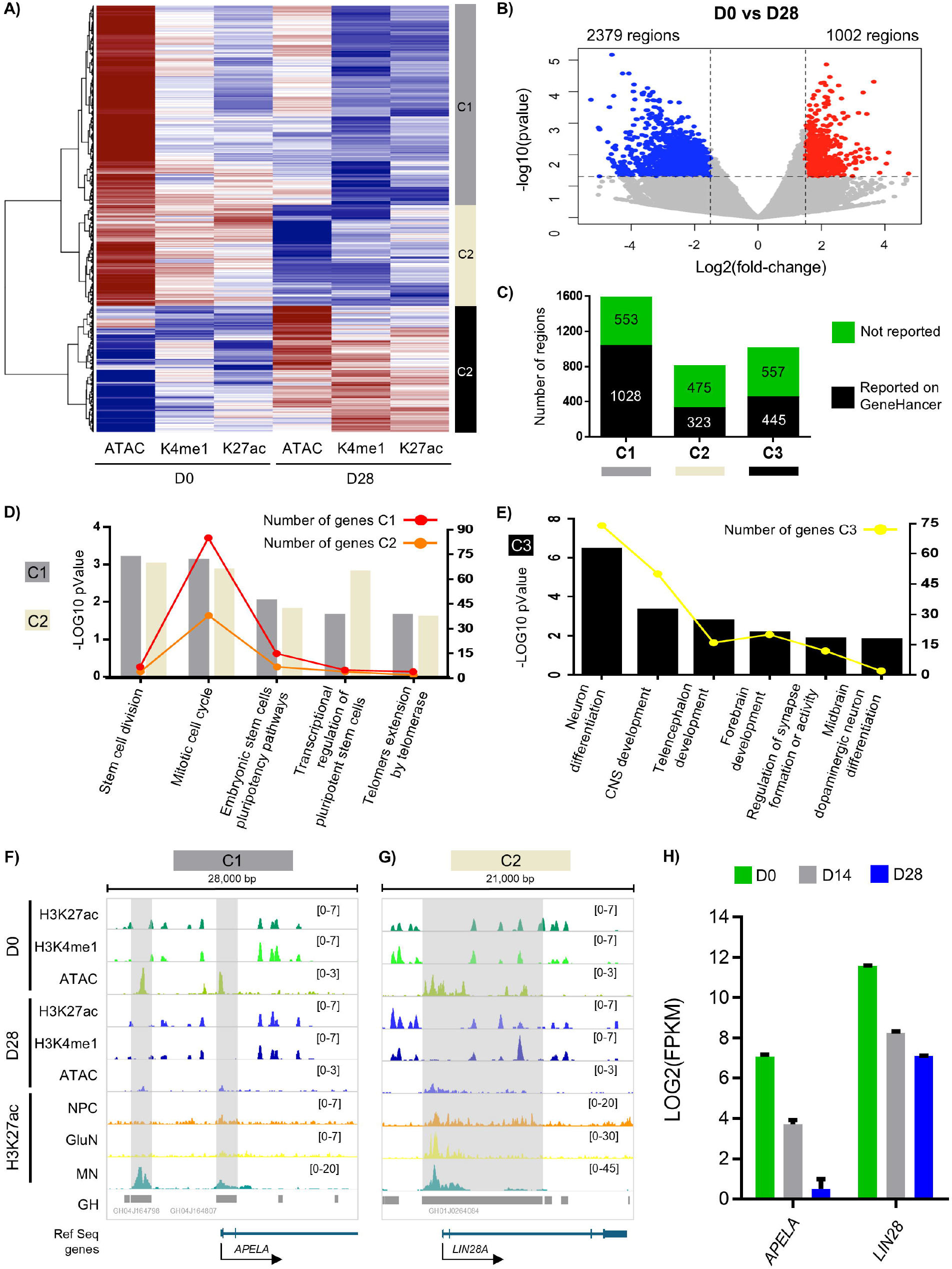
Functional annotation of active enhancers in D0 and D28 samples. **(A)** Heatmap showing the distribution of active enhancers across three clusters defined by ATAC-Seq, H3K4me1, and H3K27ac enrichment patterns at D0 and D28. C1: cluster 1, C2, cluster 2, C3: cluster 3. **(B)** Volcano plot displaying the same enhancer regions shown in panel A, plotted according to fold-change in chromatin accessibility and histone modification enrichment (x-axis) and statistical significance (–log□□ p-value, y-axis) between D0 and D28. **(C)** Number of regions per cluster that are reported (black) or not reported (green) in the GeneHancer database. **(D)** Plot showing the top enriched Gene Ontology terms for each cluster 1 and 2-associated gene set. Left y-axis and bars show statistical significance (–log □□ p-value); gray for cluster 1 (C1) and beige for cluster 2 (C2). Right-axis and dots show the number of genes for each category; red for cluster 1 (C1) and orange for cluster 2 (C2). **(E)** Plot showing the top enriched Gene Ontology terms for cluster 3. Left y-axis and black bars show statistical significance (–log□□ p-value). The right-axis and yellow dots show the number of genes for each category. **(F-G)** Genome browser tracks show representative regions from cluster 1 (C1, grey) and cluster 2 (C2, beige) near the APELA and LIN28A loci, respectively. Tracks display H3K27ac, H3K4me1, and ATAC-seq profiles in D0 and D28 samples, along with H3K27ac profiles from neural progenitor cells (NPC), glutamatergic neurons (GluN), motor neurons (MN), and GeneHancer annotations (GH). Signal intensity ranges are indicated in brackets to the right of each track. **(H)** Bar plot showing the expression levels (log2 FPKM) of APELA and LIN28A at D0, D14, and D28.

In contrast, an enhancer region located approximately 7.5 kb from FOXA1’s TTS depicts chromatin accessibility and H3K27ac enrichment in both time points however, it has an increased enrichment of H3K4me1 histone modification in D28 (Figure 2B). This enhancer region is also overlapping with a GeneHancer location (GH14J037582) whose highest score target gene is FOXA1. Interestingly, both NANOG and FOXA1 genes are differentially expressed in the same time points characterized by enrichment of both histone modifications and chromatin accessibility, namely D0 and D28 respectively (Figure 2D).

To address enhancers potentially regulating the expression of TH, a gene encoding a critical enzyme in dopamine synthesis in dopaminergic neurons, we studied both cis and distal regulatory regions surrounding TH locus. We observed that an open chromatin region located 40,742 bps upstream from TH’s TSS depicted an increased enrichment of H3K4me1 mark in D28 whereas H3K27ac modification was constant in both time points (Figure 2C). Interestingly, this region was also identified in GeneHancer database (GH11J002212) with TH as its target gene. Moreover, TH gene is differentially expressed from D0 to D28 samples (Figure 2D). These results suggest that this region is an active enhancer that gains additional H3K4me1 enrichment at D28 potentially reinforcing its enhancer activity.

### Comparison of D28 H3K27ac profiles to ENCODE related cell types

Several large-scale consortia have profiled the epigenomes of various cell-types including ENCODE (26), which analyzed chromatin states in cell lines, in vitro differentiated cells, organoids, primary cells, and tissues. To determine whether the H3K27ac profile of our D28 sample is comparable to related cell types included in the ENCODE reference epigenomes, we downloaded H3K27ac ChIP-seq datasets corresponding to NPC and motor neurons. Additionally, we also downloaded from GEO glutamatergic and GABAaergic H3K27ac profiles derived through Cut&Run assays. We calculated the Jaccard similarity coefficient to compare H3K27ac profiles across cell types in a pairwise manner. D28 H3K27ac profiles were more similar to NPC and motor neuron samples than to highly specialized glutamatergic and GABAergic neurons. Moreover, we observed a higher similarity of motor neuron H3K27ac profiles with those of NPC, glutamatergic, and GABAergic cells compared to D28 cells, potentially characterizing their neuronal lineage.

### Active enhancers depict dynamic profiles during dopaminergic induction

To study the dynamics of active enhancers in the transition from hESC to early mDAs, we generated a set of consensus regions (n=324,712) by merging significant peaks found in both time points representing histone modifications and chromatin accessibility. Next, for ChIP-seq samples (D0 and D28, H3K27ac and H3K4me1) and ATAC-seq data sets (D0 and D28) we calculated the number of counts in each consensus region. We normalized counts and adjusted for modification type (H3K27ac, H3K4me1, and ATAC-seq) and time point (D0 and D28), to perform a differential enrichment analysis using DESeq2 v1.38.3 (16). We found 3,381 differentially enriched regions with log2(fold-change) > 1.5 and p-value < 0.05. To identify dynamic enrichment profiles, we performed hierarchical clustering of differentially enriched regions and found 3 distinct clusters (Figure 3A). Regions in cluster 1 (n=1,581, gray color) depict high chromatin accessibility and a relatively low enrichment of H3K4me1 and H3K27ac marks in D0 whereas both chromatin accessibility and histone enrichments are decreased in D28. The profiles of cluster 2 (n=798, white bone color) are similar to those of cluster 1, however the histone enrichments at D0 are higher and thus decrease more drastically in D28. Furthermore, chromatin accessibility in D28 of this cluster is lost, rendering these regions potentially inaccessible for transcription factor binding. In contrast, regions of cluster 3 (n=1,002, black color) gain accessibility and enrichment of both histone modifications from D0 to D28. These results suggest that regions in clusters 1 and 2 represent enhancers active in D0 whereas enhancers in cluster 3 are active in D28. As observed in Figure 3B, a total of 2,379 and 1,002 regions depicted a decreased and increased significant enrichment of histone/accessibility marks respectively from D0 to D28.

Interestingly, we found that 63%, 38%, and 43% of the regions assigned to clusters 1, 2, and 3 correspondingly regions annotated as enhancers in GeneHancer database (17) (Figure 3C). Differentially expressed regions, clusters and overlapping GeneHancer regions are included in Additional File 4.

### Active enhancers are associated with pluripotency or neuronal differentiation pathways in D0 and D28

To infer the functions of active enhancers in each cluster, we performed a gene-set enrichment analysis using only differentially enriched regions that overlapped GeneHancer elements. GeneHancer regions are annotated with several target genes ranging from 1 to 101, demonstrating that enhancers potentially regulate multiple genes. For each region overlapping a GeneHancer element, we selected target genes which were either expressed in D0 or D28 with mean FPKM > 3 according to previously published RNA-seq data (13). We used the list of expressed target genes and performed a gene-set enrichment analysis (GSEA) for each cluster of regions using a hypergeometric statistical test. We used collections of human sets obtained from MSigDB including Hallmark gene-sets, Biocarta, Reactome, KEGG, Wikipathways, and Gene ontologies (27) (28). Gene-sets were considered significantly enriched with p-value < 0.05 and at least two overlapping genes. We found 4456, 2460, and 2154 gene-sets enriched with expressed target genes corresponding to clusters 1, 2, and 3 respectively. Among the enriched gene-sets of clusters 1 and 2, depicting targets of active enhancers in D0, we found pathways related to stem cell division and cell cycle, pluripotency, and extension of telomeres (Figure 3D). However, numerous other pathways are also significantly enriched, demonstrating that D0 cells are pluripotent and can adopt other lineages. In contrast, genes in cluster 3, depicting targets of active enhancers in D28, are enriched in pathways including neurogenesis and neuron differentiation, telencephalon and forebrain development, synapse formation and activity, and midbrain dopaminergic neuron differentiation (Figure 3E). The list of enriched gene-sets from each cluster are contained in Additional File 5.

### Active enhancers correlate with their targets’ differential expression between D0 and D28

To compare the enrichment of regions defined as active enhancers (histone enrichment + chromatin accessibility) in each time point with the expression fold-change of genes potentially regulated by these active enhancers, we annotated the 3,381 differentially enriched enhancer regions with their potential targets. Enhancer regions found in GeneHancer database were associated to their targets, however, since GeneHancer elements may have multiple targets, we selected those which were expressed FPKM > 3 either in D0 or D28 according to previously published profiles (13). Regions not found in GeneHancer database were associated to their closest gene neighbors. Next, we compared and correlated the log2-transformed fold-changes of signal enrichment (histones + chromatin accessibility) and targets’ gene expression log2 fold change (Additional File 6) and found that even though the Pearson correlation is significant (p-value < 0.05), it is positive and low (ρ = 0.16). Interestingly, by including in the comparison only regions with target genes depicting higher expression fold-changes (e.g. log2(FC) > 4), we obtain higher and significant Pearson correlation coefficients, as depicted in Supplementary figure 1.

Among the top 10 target genes with the highest log2-transformed fold-change, and significantly enriched enhancer marks, we found DLK1, RFX4, APELA, TDGF1, TAGLN3, LEFTY1, FA2H, MAP2, RTL1, and BANCR. Most enhancer-target pairs depict the same direction of fold-change, as observed by the projected linear regression in Figure 4A. Analyzing specific examples, we observe two active enhancers overlapping GeneHancer regions (GH04J164798, GH04J164807) appear to regulate the transcription of APELA. Both active enhancer regions are characterized by a decreased enrichment of chromatin accessibility and stable H3K4me1 and H3K27ac histone modifications. Interestingly, H3K27ac enrichment is high in motor neurons, whereas it is absent in glutamatergic and gabaergic neurons. The decreased active enhancer marks in these two regions are correlated to a decrease in APELA’s gene expression, suggesting a potential regulatory mechanism.

**Figure 4.**
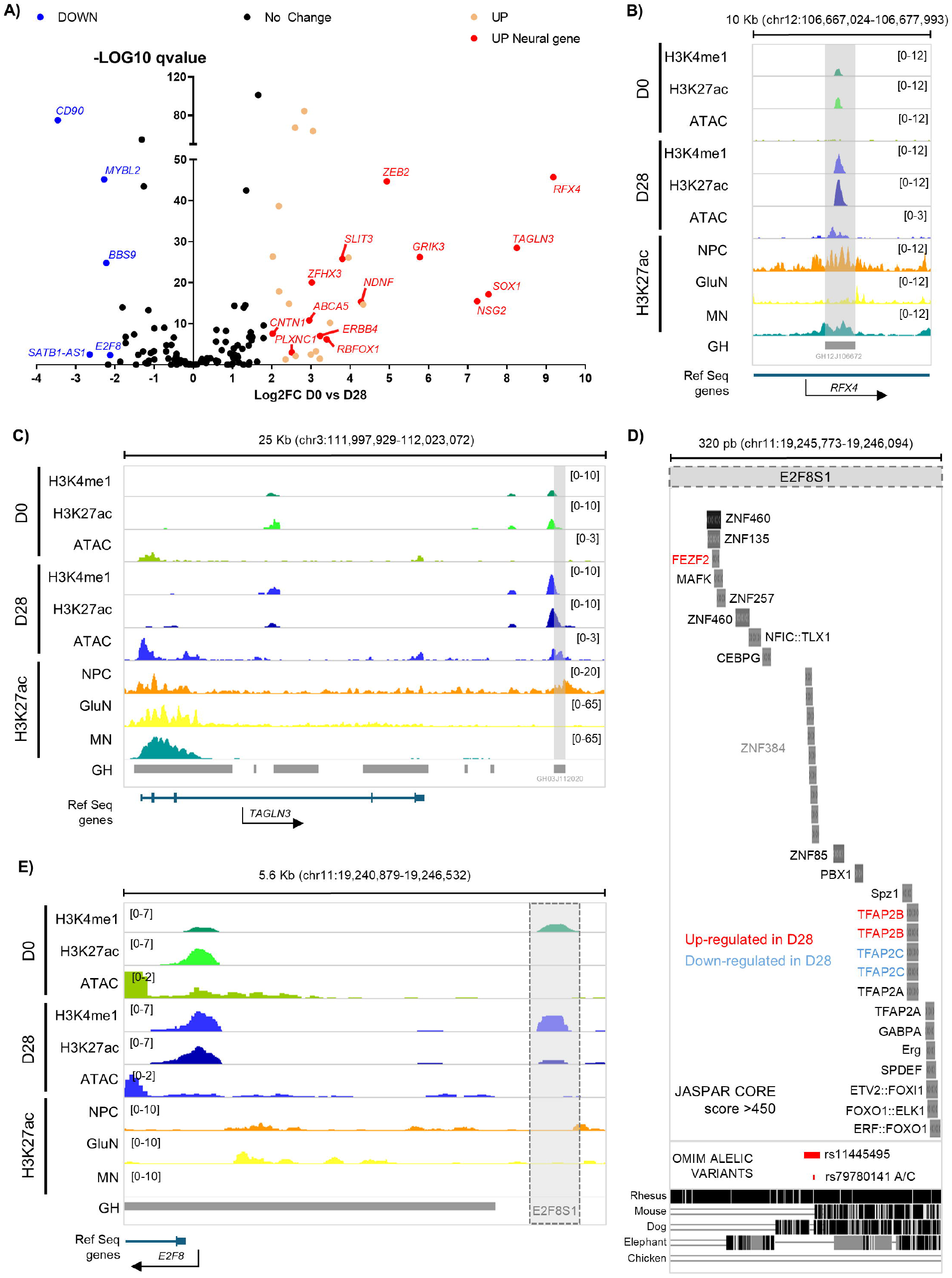
Putative enhancers in cluster 3 are linked to neuronal gene upregulation during differentiation. **(A)** Volcano plot showing differential expression of genes connected through GeneHancer annotations to cluster 3 (C3) regions (as defined in Figure 3). The x-axis shows the log2 fold change (D0 vs. D28), and the y-axis shows the –log10 q-value. Genes with significantly decreased expression at D28 are shown in blue (DOWN), those with increased expression in beige (UP), and neural-related genes with increased expression are highlighted in red (UP Neural gene). Black dots indicate genes with no significant change. **(B-D)** Genome browser tracks showing representative active enhancer regions in D28 samples near the RFX4, TAGLN3, and E2F2 loci. Tracks display H3K27ac, H3K4me1, and ATAC-seq profiles in D0 and D28, together with H3K27ac profiles from neural progenitor cells (NPC), glutamatergic neurons (GluN), motor neurons (MN), and GeneHancer annotations (GH). **(D)** Genome browser track showing a zoom-in of the E2F8S1 region from panel **D**. Predicted binding sites of TFs with a JASPAR CORE >400 are shown. In red, TFs up-regulated in D28. In blue, TF was down-regulated in D28. In black, TF with no significant change. OMIN allelic variants and conservation tracks for Rhesus, Mouse, Dog, Elephant, and Chicken are shown.

Another compelling example is an intronic active enhancer (GH12J106672) found within RFX4 (Figure 4B). This region depicts a robust increase in both H3K4me1 and H3K27ac histone modifications as well as chromatin accessibility from D0 to D28. Furthermore, RFX4 has a log2-transformed expression fold-change of 9.15 strongly suggesting that it is regulated by this intronic region. Similarly, TAGLN3 is a target gene of an active enhancer region (GH03J112020, 10Kb downstream of gene’s TTS) whose histone enrichment and chromatin accessibility are also increased in D28, correlating with a log2 expression fold change of 8.22 (Figure 4C).

### Transcription factor binding at active enhancers

Transcription factor binding occurs at gene enhancers to regulate gene expression. For the regions of each cluster defined, we performed a motif analysis to determine transcription factor binding sites at defined active enhancers. After filtering transcription factors with a binding frequency lower than 0.1% in the enhancer regions of all three clusters, we performed a heatmap of the binding frequency of the remaining 111 transcription factors in each cluster. Interestingly, some transcription factors depict binding only in specific clusters. For example, the family of RFX transcription factors (RFX2, RFX3, and RFX4) has binding sites only in active enhancers found in Cluster 3, which correlates with higher enrichment in D28. Furthermore, these transcription factors are differentially expressed in D28 (Figure 5A and 5B).

**Figure 5.**
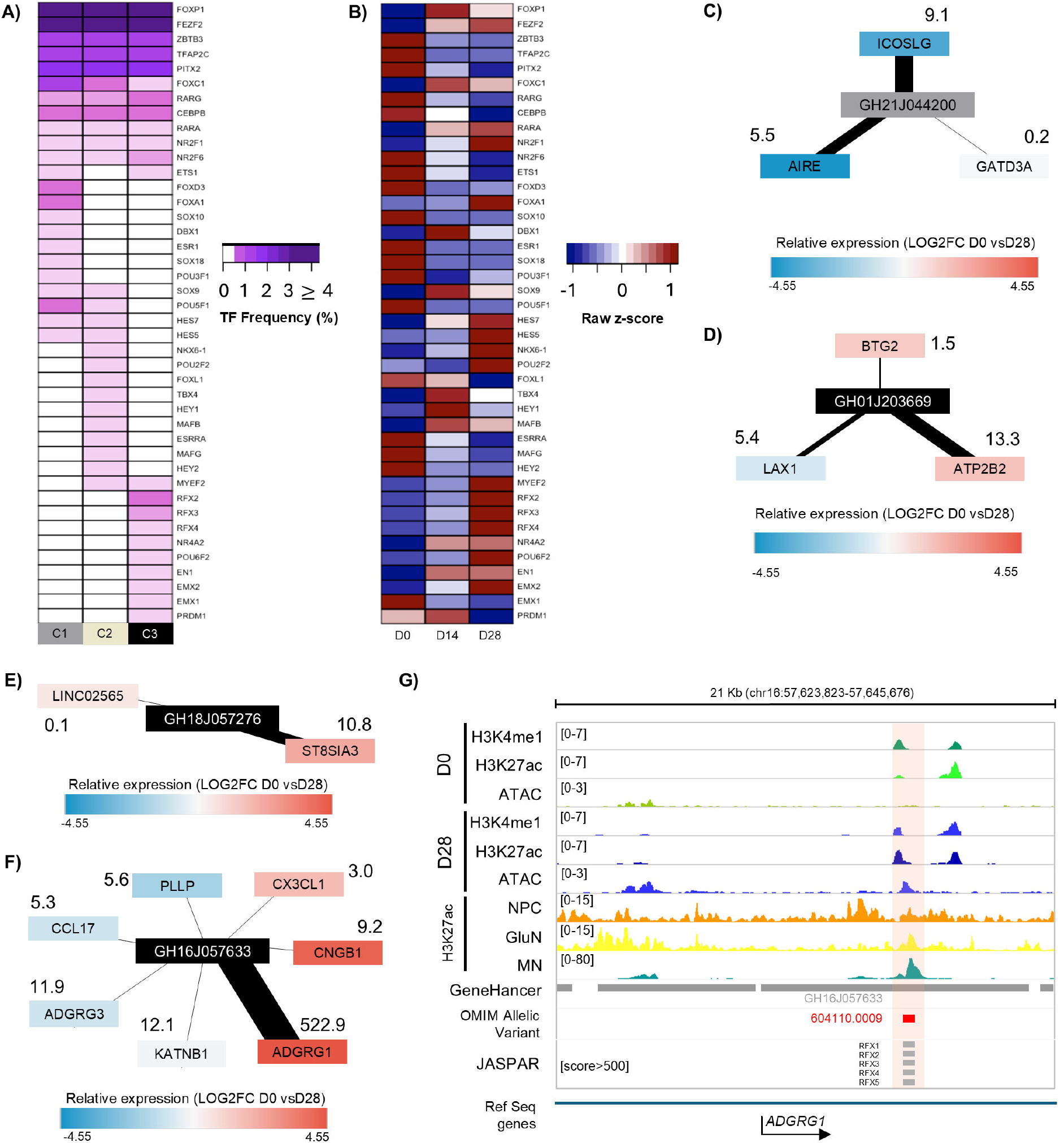
Cluster-specific transcription factor motifs and GeneHancer-linked enhancers reveal regulatory programs driving neuronal differentiation. **(A)** Enrichment of transcription factor binding motifs within clusters C1–C3, defined by regions marked with ATAC-seq accessibility and histone modifications H3K27ac and H3K4me1 (**Figure 3A**). Motifs are ranked by frequency (%) within each cluster. **(B)** RNA-seq expression dynamics of the transcription factors identified in **(A)** across dopaminergic differentiation stages (D0, D14, D28), shown as raw z-scores to highlight temporal changes in transcription factor abundance. **(C–F)** Examples of GeneHancer-annotated elements from clusters C1 in gray **(C)** and C3 in **(D-F)** with their predicted target genes. Target genes are colored by RNA-seq expression change from D0 to D28 (blue, downregulated; red, upregulated), with intensity reflecting the log2 fold change. Numbers next to each gene indicate the GeneHancer association score, and edge thickness is proportional to this score. **(G)** Genomic tracks showing H3K4me1, H3K27ac, and ATAC-seq profiles in D0 and D28 cells, as well as H3K27ac profiles in neural progenitor cells (NPC), glutamatergic neurons (GluN), and motor neurons (MN), over a 21 kb region containing the ADGRG1 locus. The highlighted enhancer (GeneHancer ID: GH16J057633) overlaps an RFX4 binding motif (JASPAR) and contains an OMIM-listed allelic variant. The enhancer belongs to the C3 region shown in panel **F**. Signal range for each track is indicated in brackets.

To further assess the correlation between differentially enriched active enhancer regions and their target genes’ expression, we built active enhancer–target networks including relative gene expression (color code). We found that active enhancer regions interact with differentially expressed genes, suggesting transcriptional regulatory mechanisms (Figure 5C-5F). We selected the GeneHancer region (GH16J057633) and its target ADGRG1 with highest prediction score (522.9). The intronic enhancer region is enriched in histone marks and chromatin accessibility in D28. Furthermore, this gene is differentially over expressed in D28 and its intronic enhancer region depicts binding sites for all transcription factors of RFX family (Figure 5G), whose members are also over expressed in D28. These results suggest that RFX4 plays an important role in dopaminergic induction through enhancer regions which become active only in D28.

## Discusion

The cellular composition of the developing human brain is highly diverse and dynamic. Over 100 different neural and non-neural phenotypes have been described using multi-omic approaches across developmental stages (29) and adulthood (30). Understanding the fine gene-regulatory mechanisms behind the complex developmental pathways involved in the induction of a specific neuronal subtype, such as the midbrain dopaminergic neurons, is crucial for elucidating brain function and some aspects of neurological diseases. Here, we focused on identifying active transcriptional enhancers on developing dopaminergic neurons by mapping active-enhancer-associated histone modifications. Our multi-omic analysis allowed the identification of 55,269 enhancers in ESCs and 13,102 in mDAs, of which 64% and 52% had not been previously annotated on the GeneHancer database. Accurate identification of enhancers’ target genes remains to be done and can be achieved by generating mDA-specific enhancer-promoter interactome maps through high resolution Hi-C (31). Additionally, ChIA-PET, HiChIP and PLAC-Seq combine techniques from biochemical annotation and gene-enhancer interaction identification to provide a fuller picture.

We also found distinct sets of transcription factors (TF) binding motifs differentially enriched between the putative enhancers of these two cellular types (ESC y mNDA), suggesting a complex interplay of enhancer activity, TF differential binding and TF differential expression in both processes: pluripotency maintenance and dopaminergic induction. We pointed out to novel TF regulating some aspects of midbrain dopaminergic induction such as MYEF2 and the RFX family. Overexpression of the RFX4 TF has been previously reported by our group and others in mammalian models of neural (32) and dopaminergic induction (13). TF motif analysis showed that D28-enriched putative enhancers frequently contain RFX4 binding motifs, in contrast with D0-enriched putative enhancers, which correlates with RFX4 transcription levels. Some of the RFX4 binding motif-containing putative enhancers have a regulatory association with neural-development genes. Accurately mapping RFX4 binding sites during dopaminergic induction is necessary to fully establish RFX4 downstream regulatory targets and its role in dopaminergic neuron biology.

We previously reported that the transcriptional changes associated with dopaminergic differentiation from hESC involves both up and down-regulation of several genes (13). There is evidence suggesting that transcriptional enhancers can act as silencer elements under specific cellular and developmental contexts (33). Some D28 specifically enriched putative regulatory elements were located in loci of downregulated genes, indicating possible silencer activity of these elements.

We found that our putative silencers, as well as many putative enhancers, were enriched in the well-known neural-specific TF FEZF2. FEZF2 can act as an activator TF that regulates multilineage neuronal differentiation (34) and has been reported to be necessary for dopaminergic neuron development (35). FEZF2 its also a well know repressor of WNT-signaling inhibitors, thus activating WNT/β-catetin signaling (36), which is necessary for DA induction (37). Furthermore, detailed characterization of this TF has shown that its regulatory roles are associated with the switch between quiescent and proliferative states of brain cells (38). Accordingly, some FEZF2-bound putative regulatory elements are located in cell-cycle regulatory genes loci, such as E2F8 (39). FEZF2 bound targets during dopaminergic induction and patterning remain to be experimentally assessed to fully understand this TF role in cell cycle exit regulation and mNDA maturation.

## Supporting information

Additional file 1 - Peaks

Additional file 2 - Peak annotation

Additional file 3 - overlapping histones atac GH

Additional File 5 - GSEA

Additional File 6 - DEGs targets enrichment expression

Additional File 7 motifAnalysis

Additional File 4 - norm counts DEGs

## Ethics approval and consent to participate

Not applicable

## Consent for publication

Not applicable

## Authors’ contributions

R.CD-D, I.V. and E.S-R supervised the project. C.M-R and M.G-L designed the project, performed experiments and analysis, and wrote the manuscript. R.CD-D performed analysis and wrote the manuscript. A.D-M, E.S-R and R.G.R-V performed experiments and analysis. A.L-O performed experiments. All authors reviewed and approved the manuscript.

